# Phase and amplitude correlations change with disease progression in idiopathic Rapid eye-movement sleep behavior disorder patients

**DOI:** 10.1101/2021.03.12.435081

**Authors:** M. Roascio, A. Canessa, R.D. Tro, P. Mattioli, F. Famà, L. Giorgetti, N. Girtler, B. Orso, S. Morbelli, F. M. Nobili, D. Arnaldi, G. Arnulfo

## Abstract

Cognitive decline is a common trait of neurodegenerative diseases of central nervous system and one of the major risk factors associated with faster phenoconversion from prodromal stages. In the transition to full-blown clinical syndromes, increased phase synchronization in the theta, alpha or beta EEG rhythms is thought to reflect the activation of compensatory mechanisms that may counterbalance the cognitive decline of patients affected by Mild Cognitive Impairment (MCI). Patients suffering from idiopathic Rapid eye-movement sleep Behavior Disorder (iRBD) have high risk of developing Parkinson Disease (PD) or Dementia with Lewy Bodies (DLB) and cognitive impairment is among the strongest risk factors together with motor symptoms. Here we wanted to investigate whether altered phase synchronization and amplitude couplings of the brain oscillations could be linked to the balancing of cognitive decline in a longitudinal cohort (N=18) of iRBD patients. We measured high-density Electroencephalographic (HD-EEG) activity at baseline and follow-up and quantified power distribution, orthogonalized amplitude correlation and weighted phase lag index. Despite the overt neurodegenerative progression (three patients converted to PD and one to DLB), cognitive decline was not evident from Mini Mental State Examination (MMSE) or neuropsychological tests. On the other hand, alpha phase synchronization and delta amplitude correlations were significantly different at follow-up compared to baseline. In particular, alpha synchrony was enhanced while delta amplitude coupling was reduced. Those differences were more pronounced among central-posterior channels while frontal channels showed a reduced number of significant edges with respect to surrogates. Both large-scale amplitude and phase coupling significantly correlated with cognitive or neuropsychological scores but not with sleep quality indices. Altogether, these results suggest that increased alpha phase-synchronization and reduced delta amplitude correlation may be considered as electrophysiological signs of an active compensatory mechanism of the cognitive impairment in RDB patients. Large-scale functional modifications could thus be used as significant biomarker in the characterization of prodromal stages of PD.

**Statement of Significance:** Cognitive impairment and RBD emerge much earlier than the better-known motor symptoms distinctive of synucleinopathies. An improved investigation of RBD may constitute an important biomarker for an early diagnosis of the actual neurodegenerative diseases. For the first time, this preliminary study aims to quantify the large-scale network couplings as electrophysiological manifestation of the compensatory mechanism to the cognitive impairment in a longitudinal study of idiopathic RBD patients. Unfortunately, the small number of the subjects limits the generalizability of our observations, but this is only preliminary works in a larger project that aims to investigate advanced electrophysiological markers for an early diagnosis of the synucleinopathies.

## 1. Introduction

Rapid eye movement sleep Behavior Disorder (RBD) is a parasomnia that involves violent and undesirable behaviors^1^ such as the physical reaction to dream due to the loss of normal muscle atonia during rapid eye movement (REM) sleep. RBD can occur alone (idiopathic RBD – iRBD) or in association with other neurological disorders (symptomatic RBD)^2^ and iRBD patients have high risk of developing Parkinson’s Disease (PD), Dementia with Lewy Bodies (DLB) and multiple systemic atrophy.^3,4^

Clinical RBD symptoms include cognitive decline,^4^ impaired olfaction and color discrimination,^5^ abnormal metabolic network activity^6^ and reduced striatal dopamine transporter (DAT) uptake.^3^

Compared to healthy subjects, RBD patients show a reduction of dominant alpha rhythms in occipital, parietal and temporal lobes^7^ along with a power increase in delta and theta frequency range in frontal and central lobes.^8–10^ On the other hands, in magnetoencephalographic (MEG) studies, the increase of the theta, alpha and beta phase synchronization have been associated with a compensatory mechanism to the cognitive decline in Mild-Cognitive Impairment (MCI).^11,12^ In MCI, the hyper-synchrony in theta and beta band have been also interpret as a sign of the neurodegenerative process where a loss of inhibitory neurons is caused by the amyloid toxicity.^13^ We hypothesized that hyper-synchronization and altered coupling could be a common feature in prodromal stages of several neurodegenerative diseases. However, it remains unclear how such electrophysiological biomarkers evolve over time in prodromal synucleinopathies. Here, we used a high-density EEG (HD-EEG) system to investigate the disease progression-related evolution in the large-scale brain network couplings in a group of iRBD patients and we correlated finding with longitudinal cognitive and nigro-striatal dopaminergic single photon emission tomography (SPECT) data.

## 2. Materials and Methods

### 2.1. Data acquisition

A total of 22 iRBD patients (21 men; mean age 70 ± 6.8 years at the first clinical evaluation) were recruited at sleep outpatient facility of University Neurology Clinics at Policlinico San Martino in Genoa and they were investigated in two subsequent sessions, namely at baseline and at follow-up (after 24.2 ± 5.9 months; range: 14 – 41 months) including clinical, neuropsychological, EEG and DAT-SPECT assessment. The diagnosis of idiopathic RBD was made according to international criteria (ICSD 3) and was confirmed by overnight video-polysomnography. In accordance with the declaration of Helsinki, all participants gave informed consent before entering the study, which was approved by the local ethics committee.

#### 2.1.1. Clinical assessment

The patients underwent general and neurological examination to exclude other neurological and psychiatric disorders. Brain Magnetic Resonance Imaging (MRI), or computed tomography in the case MRI was unfeasible, was used to rule out brain diseases such as tumors or lesions. The presence of white matter hyper-intensities was not an exclusion criteria if the Wahlund scale was not >1 for each brain region.^14^ Patients underwent baseline clinical evaluation, including: i) the Mini-Mental State Examination (MMSE) as a global measure of cognitive impairment; ii) the Movement Disorder Society-sponsored revision of the unified Parkinson’s disease rating scale, motor section (MDS-UPDRS-III) to evaluate the presence of parkinsonian signs; ii) clinical interview and questionnaires for activities of daily living (ADL) and instrumental ADL to exclude dementia; iii) Beck depression inventory II (BDI-II) to rate depressive symptoms; iv) the Italian version of Parkinson Disease Sleep Scale version 2 (PDSS-2) was used as a clinical measure of sleep disorders.^15^

#### 2.1.2. Neuropsychological assessment

Patients underwent a comprehensive neuropsychological (NPS) assessment, evaluating the five main neuropsychological domains: language, executive functions, visuospatial abilities, memory, attention and working memory. We administered the following neuropsychological tests: semantic verbal fluency and phonemic verbal fluency, Stroop color word, Stroop color, Trail making test A and B (TMA, TMB), clock completion, constructional apraxia, simple copy and copy with guiding landmarks, Rey Auditory Verbal Memory Test (RAVLT, immediate and delayed recall), Babcock story, Corsi span, digit span and symbol digit. References and normative data are detailed in a previous study.^16^ In the identification the variables expressing a similar part of total variance of the baseline native neuropsychological measures, we used a factor analysis with varimax rotation to minimize multicollinearity and to reduce the number of neuropsychological variables. We set a conventional threshold of 0.4 to factor loadings (expressing the factor-variable correlation) to select the group of variables mainly represented by each factor. Factor analysis identified four principal factors (Tab. S1). Factor one was mainly related to visuospatial abilities (NPS-VS), factor two to verbal memory (NPS-VM), factor three to executive functions (NPS-EX), and factor four to attention (NPS-AT), respectively.

#### 2.1.3. Electrophysiological signal recording

All iRBD patients underwent HD-EEG evaluation during relaxed wakefulness within three months from diagnosis, late in the morning to minimize drowsiness. For each session, the acquisition protocol consisted in about 22.57±2.62 minutes (range: min 17, max 29) of resting-state subdivided in: 1.03±0.67 minutes (range: min 0, max 3) with eyes-open, 3.8±0.97 minutes (range: min 0, max 5) during hyperventilation and 17.75±2.93 minutes (range: min 12, max 26) minutes with eyes-closed. We used the Galileo system (EBNeuro, Florence, IT) to acquire bandpassed (0.3 – 100 Hz) signals from 64 electrodes, at a sampling rate of 512 Hz. Electrodes were placed according to the 10-10 International System where the reference electrode and ground were Fpz and Oz, respectively (Fig. 1). We recorded simultaneously the horizontal electro-oculogram to monitor eye movements with the same recording parameters of EEG. Electrode impedances were closely monitored and kept below 5 kOhm. An EEG technician monitored the recording session to maintain a constant level of vigilance of the patient, prevent sleep and to preserve a high signal quality across the whole recording session. For each patient and for both visits, we selected the eyes-closed condition for further analyses.

**Figure 1:**
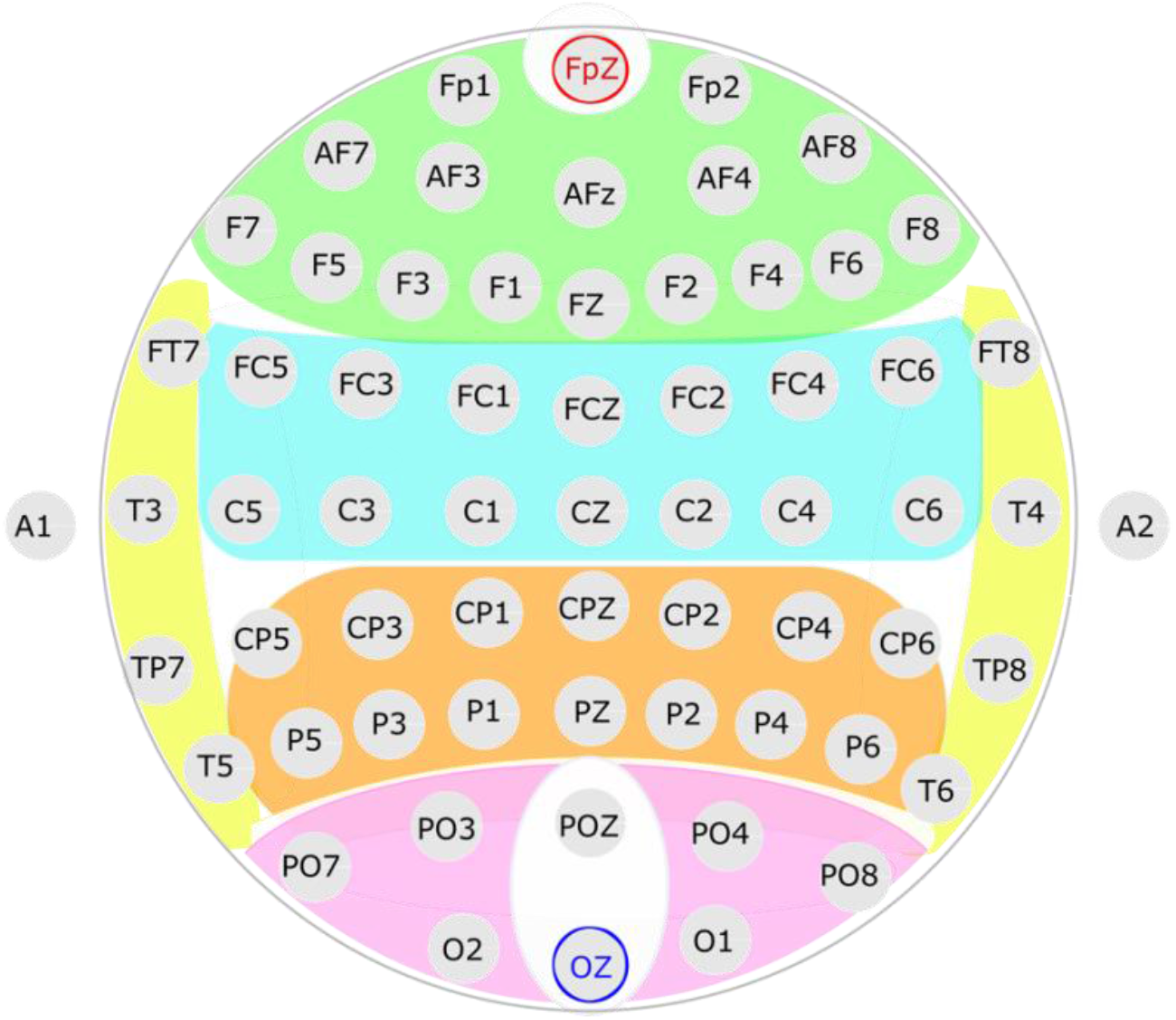
Electrode channel groups. Groups of EEG sensors divided in 5 structural area: frontal (green), central (azure), parietal (orange), occipital (pink) and temporal (yellow). Fpz (red) and Oz (blue) are the reference electrode and ground, respectively.

As control dataset, we selected 10 healthy subjects from Mind-Brain-Body dataset of the Max Plank Institute.^17^ The EEG data were acquired using BrainAmp MR plus amplifier using 61-channel active ActiCAP electrodes positioned according to 10-10 system. The EEG data consist of 8 minutes of eyes-closed resting-state condition.

#### 2.1.4. Molecular imaging evaluation

Within three months from diagnosis, all iRBD patients underwent [^123^I]N-ω-fluoropropyl-2β- carbomethoxy - 3β -(4-iodophenyl)nortropane (FP-CIT) SPECT to measure the striatal dopamine reuptake transporters (DAT) density according to EANM guidelines.^18^ Details on SPECT data acquisition and analysis could be found in supplementary materials. Reconstructed images were exported in the Analyze file format and processed by the Basal Ganglia V2 software^19^ to compute specific to non-displaceable binding ratio (SBRs). In particular, background uptake (occipital region) was subtracted by putamen or caudate uptake as follows: (putamen/caudate uptake – background uptake)/background uptake, to compute SBRs values. For subsequent analyses, we computed a mean SBR values between right and left hemisphere for both caudate and putamen.

### 2.2. Data pre-processing

Data pre-processing and analysis was carried out in MATLAB, R2019a using Brainstorm^20^ and custom scripts.

As a first step, we preprocessed EEG data of iRBD patients. We used a band pass finite impulse response (FIR) filter (1-80 Hz, Kaiser window, order 1858) to eliminate the low and high frequency artefacts and we applied zero-phase infinite impulse response (IIR) notch filter (order 2) to remove the power line noise (50 Hz).

Later, we visually removed all channels and time windows showing artefactual activity such as blinks, muscular activity and other sporadic artefacts. We removed an average of 19 channels from each patient (range: min 3, max 38). In particular, for all subjects we rejected channels A1, A2 and POZ since the percentage of artefactual windows was higher than 90% due to bad contact between scalp and the electrode. Moreover, we rejected the time windows that showed evidence of drowsiness defined as those time intervals in which slow-wave activity replaced the occipital alpha rhythm.

Four subjects were excluded from the successive analysis due to an excessive artefactual activity, which left less than 5 minutes of pruned eye-closed resting-state data after the cleaning. The final population size for this study was hence 18 patients.

The 10 healthy control subjects dataset contained in Mind-Brain-Body dataset were already pruned and bandpass filtered (1-45 Hz, 8^th^ order, zero-phase Butterworth filter).

For all cleaned datasets, we applied the Scalp Current Density (SCD) reference montage^21,22^ to all clean sensors with spline method (lambda 0.00001, order 4, degree 14) and down-sampled the resulting time-series at 100Hz. We defined five regions of interest (ROI) for the subsequent analyses by grouping electrode contacts in frontal, central, occipital, parietal, and temporal (Fig. 1) in accordance with their position on the scalp.

### 2.3. Spectral Analysis

We computed the Power Spectral Density (PSD) for all channels and for all sessions in both the iRBD patients and healthy control subjects. The spectral analysis was performed using Welch modified periodogram method (pwelch.m - MatLab) between 1 Hz and 30 Hz, with a resolution of 1 Hz. We computed the Absolute (AP) and Relative Power (RP) for four frequency bands of interest (BOI) corresponding to common brain rhythms, *i.e.* δ (1-4 Hz), θ (5-7 Hz), α (8-13 Hz), and β (13-30 Hz).

For each BOI *b* the AP is computed as

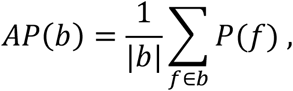

where *p*(*f*) is grand average power computed as the PSD average across all the channels as,

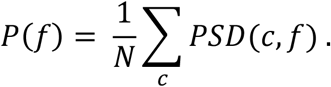

where *N* is the number of channels considered.

Once computed the AP, the RP of each BOI *b* is defined accordingly as:

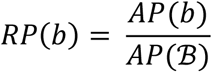

where *AP(ℬ)* is the AP computed over the whole frequency range considered *ℬ* (1-30Hz).

The simultaneous presence of an aperiodic background with *1/f* decaying spectrum superimposed to the true oscillatory periodic components characterizes the EEG recordings. The changes in the slope of the *1/f* background activity could be representative of aging and, if pronounced, suggest an on-going pathological degeneration process.^23^ To properly analyze specific power changes, we decided to parameterize the estimated PSD. In this work we modeled the PSD as the sum of putative, periodic oscillatory components parameterized by their central frequency, power and bandwidth, as measured from Gaussian mixture model fits, plus an aperiodic component, as described by a power law fit *f*^−α. 23^ Following the parameterization procedure, we computed the slope α of the aperiodic component to observe whether there was any difference between the two recording sessions.

### 2.4. Phase and amplitude correlation analyses

We computed phase-synchronization and amplitude correlations across a range of frequencies for all channel pairs and for all sessions in both iRBD patients and healthy control subjects. We analyzed the broad-band SCD-referenced time-series by means of a time-frequency decomposition using 15 narrow-band Morlet wavelets in the 1-15 Hz range with 3s time-widths at 1Hz.^24^

Volume conduction, signal mixing, and source leakage significantly inflate phase synchronization analyses of sensors EEG data^25,26^ and reduce possibilities to detect true significant changes in phase-synchronization profiles. To minimize this, we adopted the weighted Phase-Lag Index^25^ (wPLI) as it is insensitive to volume conduction and has an increased statistical power^25,26^ compared to other phase synchronization metrics.

The wPLI is defined as

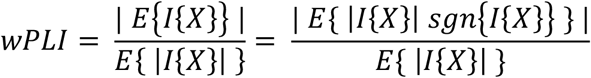

where I{x} denotes the imaginary part and *X* is the cross-spectrum between two EEG channels.

While phase synchronization reflects a mechanism of neuronal communication per se,^27^ coherent amplitude modulation between distant brain regions could reflect the concurrent activation of different neuronal population in response to the same sensory stimulation. Hence, amplitude correlation could be seen as an electrophysiological correlate of Blood Oxygenation Level Dependent (BOLD) signal. As in phase synchronization, volume conduction and source leakage mask true significant coupling. As volume conduction insensitive measure, we adopted the orthogonalized Correlation Coefficient (oCC) that is defined as the Pearson correlation coefficient between two orthogonalized time series^28^. We computed oCC as:

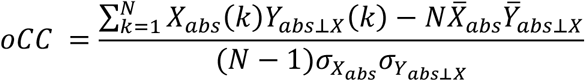

where *X_abs_* is the absolute value of the analyzed complex time series with standard deviation 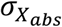 and mean value 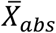, *_Yabs⊥X_* is the absolute value of the complex signal *Y(t)* orthogonalized to the complex signal *X(t)*, with its respective standard deviation 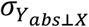and mean value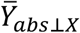, *N* denotes the time series, and *k* the samples number.

Both wPLI and oCC are limited in [0,1] range where 0 represents absence of phase-synchronization or amplitude correlation.

### 2.5. Statistical analysis

We conducted a preliminary statistical analysis to investigate the cognitive decline in iRBD patients and their neurodegeneration process. First, we applied the Kolmogorov-Smirnov test (kstest.m - MatLab) to observe whether clinical, neuropsychological and imaging assessment came from a standard normal distribution. Given the absence of normality, we used non-parametric Wilcoxon Test (ranksum.m –MatLab) to investigate if clinical, NPS and DAT-SPECT SBRs showed significant changes over time between baseline and follow-up. Later, we wanted to test for systematic differences in EEG power between the two recording sessions in iRBD patients. We thus applied a two-ways repeated measure analysis of variance (rm_anova2.m - MatLab - File Exchange)^29^ to evaluate the statistical effects of channel groups and between the two recording sessions in AP, RP and in the slope values.

In order to quantify the spatial extent of phase synchronization and amplitude couplings, we computed the fraction of significant, for both iRBD patients and healthy control subjects. The fraction of significant is defined as the ratio between the number of statistically significant channel pairs out of the total number of channel pairs. Since under null-hypothesis of no coupling both wPLI and oCC show a Gaussian distribution with mean μ and variance σ^2^ equal to the mean and the variance of the surrogates. For each pair of channel pair and a given frequency, we obtained a single surrogate by arbitrary rotating the time-series of the second channel in one pair and we computed the wPLI and oCC metric between the original signal of the first against the rotated version of second one. To identify the threshold of significance we z-scored the metrics and used z-score above 2.33 as significance threshold of 0.02.

We investigated the difference of the wPLI and the oCC between the two recording sessions of iRBD patients to assess putative disease progression effects on large-scale brain networks. We wanted to observe whether the mean difference of observed data of wPLI/oCC was statistically significant. As a first step, we thus applied a Multiple Comparison Permutation Test^30^ to calculate surrogates data in order to fix a threshold of significance for the difference.

In particular, we computed surrogate data mixing the values of observed data in the two recording sessions. We split the mixed values in two new vectors (one for each recording sessions) with the same dimensionality of two original categories. We, thus, computed the difference between the two new vectors for each subject and we repeated all this procedure 1000 times. Since during multiple permutations different data selection could yield an unbalanced variance between permutations, we applied a normalization. This standard procedure encompass the normalization for each frequency by dividing each permutation for the global standard deviation computed across permutations.^30^ Finally, we selected for each permutated values the absolute maximum of the normalized surrogates across frequency (max T-statistics Multiple Comparison Correction). We defined the upper and lower bounds of confidence limits for the difference as the 2.5 and 97.5 percentiles across permutations. The mean difference between the observed data at baseline and follow-up was considered statistically significant when it lied outside the threshold range (*p* < 0.05). Given the limited size of our cohort, we adopted a more conservative approach and we considered a significant effect on the difference between baseline and follow-up only those frequency points where both the confidence limits of the population variance (bootstrap, N=1000) were outside the threshold range.

As a last statistical analysis, we investigated whether there was a correlation between the altered phase and amplitude couplings with clinical, NPS and DAT-SPECT data in RBD patients. Hence, we applied the Partial Pearson Correlation Coefficient (partialcorr.m – MatLab) to this data controlling for the age of the patients at baseline and follow-up. Finally, we corrected the p-values with Benjamini-Hochberg (BH) method (fdr_bh.m – MatLab – File Exchange)^31^ for multiple hypothesis testing.

## 3. Results

After 24.4 ± 6.1 months of follow-up, four patients (22%) developed a neurodegenerative disease (three PD and one DLB). As preliminary results, clinical, NPS and DAT-SPECT data (Tab. S2) did not show significant difference between baseline and follow-up, suggesting that cognitive reservoir and nigro-striatal dopaminergic function remained largely unchanged in the group. This is likely due to the limited number of phenoconverted patients.

### 3.1. Effects of disease progression in amplitude spectral profiles

Here, we tested whether alpha band peak slows down as disease progresses.^7^ First, we observed that the overall power spectrum distribution of iRBD patients (at baseline and at follow-up) slows down towards lower frequency compared to healthy control subjects (Fig. 2) and, indeed, recordings at both time points show a slowing of brain activity as measured by relative power increase in theta band (5-7Hz) and a reduction in alpha (8-13Hz) and in beta (13-30Hz) band compared to healthy controls subjects.

**Figure 2:**
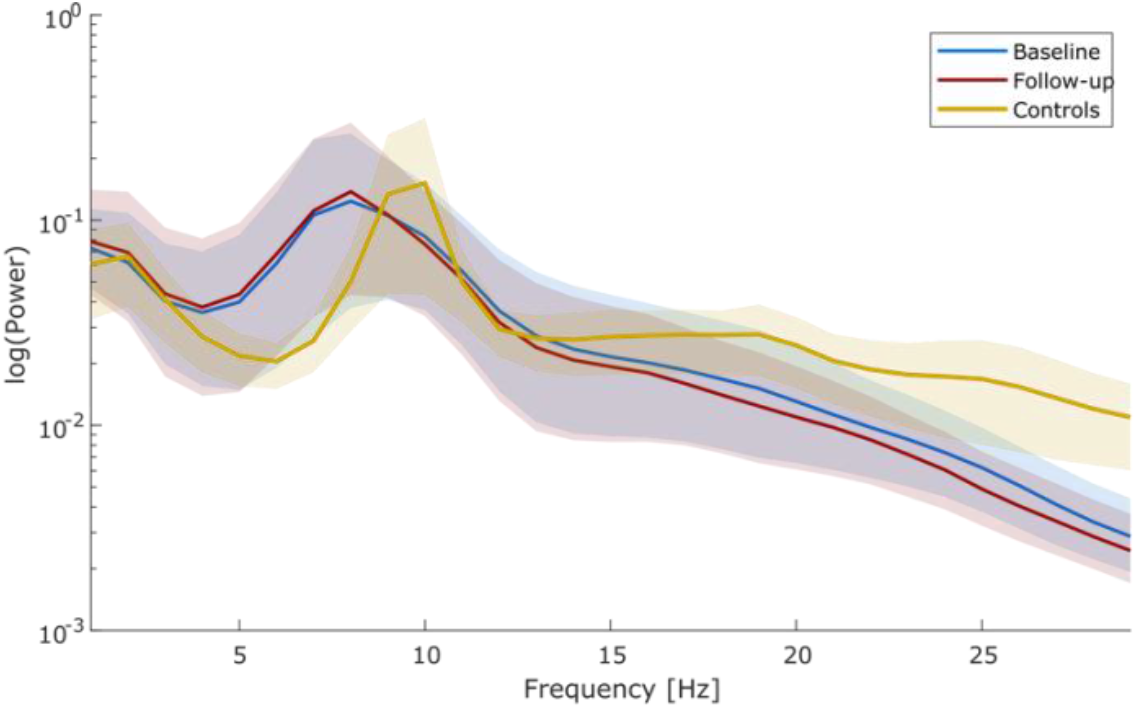
Effects of disease progression in amplitude spectral profiles. Group-level averaged power spectral densities for RBD patients at baseline (blue) and at follow-up (red), and controls group (yellow). Shaded areas represent confidence intervals at 5% around population mean. **Legend:** PSD = Power Spectral Density

Globally the PSD do not differ between the two recording sessions in iRBD patients (Fig.2). Then, we investigated for a possible effect within channel group (frontal, central, parietal, occipital and temporal). For each channel groups and each recording conditions of the iRBD patients, we evaluated the RP (Fig. S1) and the AP (Fig. S2). The RP in alpha band is significantly different across electrode groups (two-factor ANOVA, *p* = 0.0175) but not across recording sessions (Tab. S3). We reported a significant difference also in delta (*p* = 0.0106), theta (*p* = 0.0105) and alpha (*p* = 0.0017) band for the AP values (Tab. S5). All other factors and their combinations have not reached a statistical significance for AP and RP values.

Moreover, we extracted the scaling exponent of fitted power laws and evaluated whether there was a significant difference between recording sessions or EEG channels. Analogously to the results presented above, we have not observed a significant difference between recording sessions, channel locations and their interactions (Tab. S4).

Altogether, our results suggest that progressing of iRBD pathology is not associated with a significant slowing of activity at group level in the time span we collected the data.

### 3.2. Global phase and amplitude couplings change with the disease progression

We then wanted to investigate how the phase-synchronization and amplitude correlation change between baseline and follow-up.

First, we observed that both strength (Fig. 3a) and spatial extent (Fig. 3b) of global wPLI peak at the alpha band for both recording sessions and within all channel groups (Fig. S3). Furthermore, global alpha band wPLI decreases in RBD compared to controls (Fig.3a). The alpha band wPLI significantly (*p* < 0.05, permutation test – max T. corrected) increases in RBD patients at the second visit (Fig. 3c) and within all channel groups except the temporal channel group (Fig. S3). On the other hand, the large-scale amplitude coupling strength (Fig. 3d), but not spatial extent (Fig. 3e), is significantly (*p* < 0.05, permutation test – max T. corrected) reduced from the first to the second recording session in the delta band (Fig 3f) and in all channel groups except the occipital and temporal channel groups (Fig. S4). Altogether, these results highlight the global functional modifications associated with disease progression in iRBD patients.

**Figure 3:**
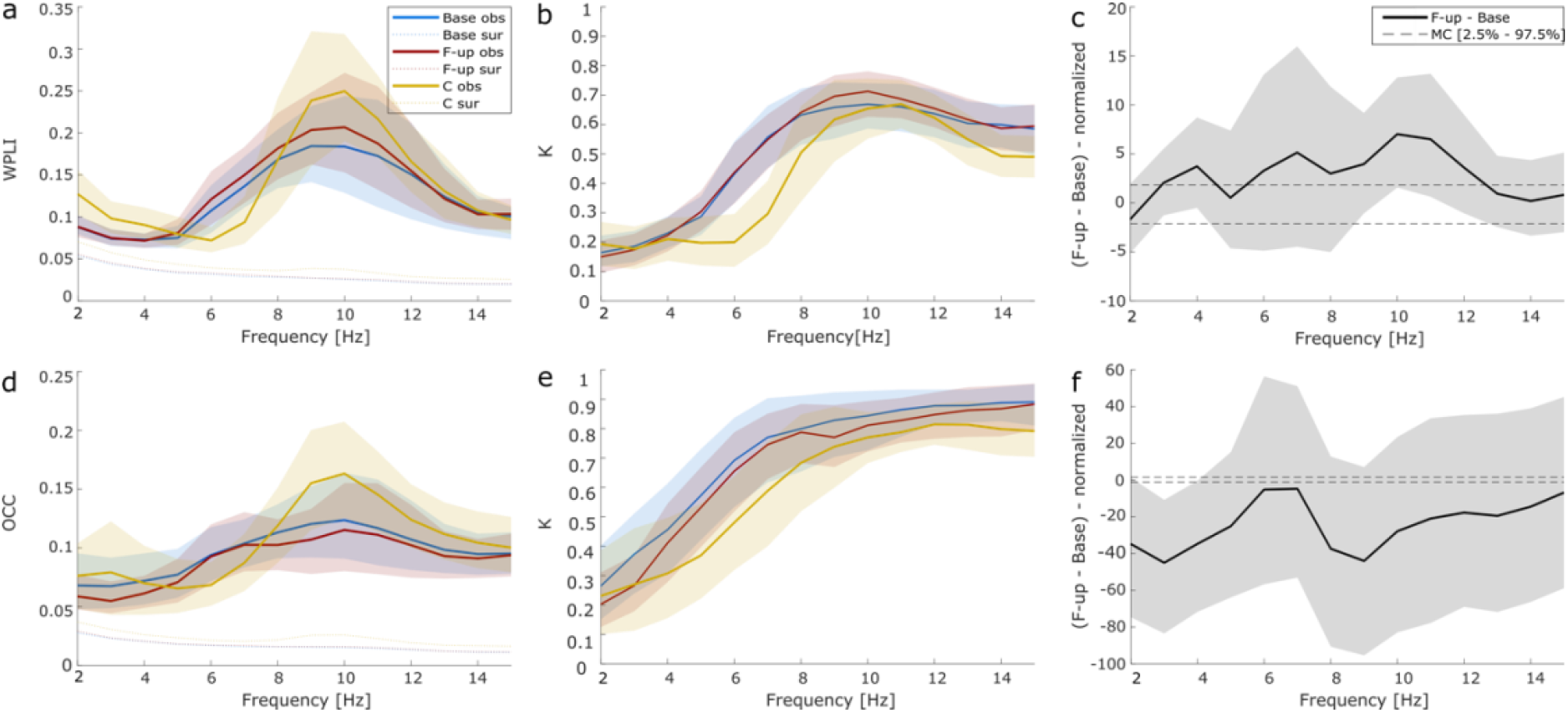
Global phase and amplitude couplings change with the disease progression. (a,d) Strength and (b, e) extent of population averaged wPLI (a,b) and oCC (d,e) at baseline (blue), at follow-up (red) and in healthy control subjects (yellow). Shaded areas represent confidence interval around population mean (1000 bootstraps). Dashed lines represent surrogate average. (c,f) Absolute difference of phase synchronization (c) and amplitude correlation (f) spectral profile between conditions. Shaded areas represent confidence intervals at 5% around mean. Dashed lines represent significance threshold for a two-tail pairwise permutation test at (p < 0.05) corrected for multiple comparison using max T. statistics across frequencies. **Legend:** wPLI = weighted Phase Lag Index; oCC = orthogonalized Correlation Coefficient; Base = baseline; F-up = follow-up; C = controls; MC = multiple comparison; k = fraction of significant.

### 3.3. Large-scale network modifications related to disease progression

We looked at the organization of the resulting large-scale brain networks and asked whether specific brain circuits were more affected as the disease progresses.

First, we modelled the brain topology as weighted graph where nodes represent EEG channels, edges are weighted proportionally to the wPLI (Fig 4a-b) and oCC values (Fig 4c-d) and specifically looked at degenerated networks in the alpha band (10Hz) and delta band (4Hz). Frontal electrode group and inter-hemispheric phase coupling showed a reduction in the number of significant (in respect to surrogates, *p* < 0.05) edges at cohort level when considering edges that were significant in at least 50% (Fig. S5 a-b), 66% (Fig. 4a-b) and 75% (Fig. S5c-d) of the subjects.

**Figure 4:**
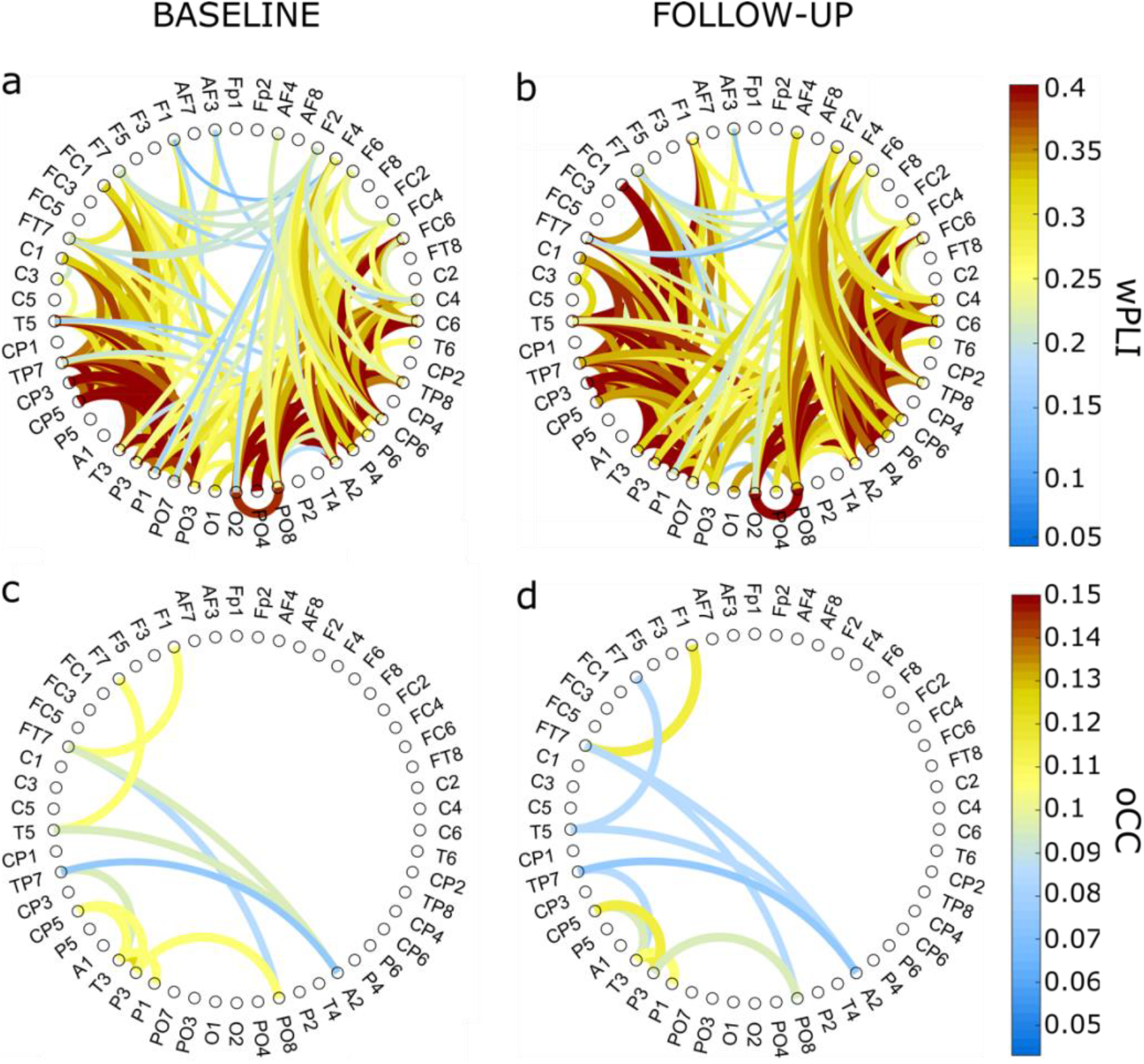
Large-scale network modifications related to disease progression. Graph of population averaged wPLI in alpha band (10Hz) at baseline (a) and at follow-up (b) for significant edges (p<0.05) in at least 66% of the patients. Graph of population averaged oCC in delta band (4Hz) at baseline (c) and at follow-up (d) for significant edges (p<0.05) in at least 66% of the patients. **Legend:** wPLI = weighted Phase Lag Index; oCC = orthogonalized Correlation Coefficient.

We then observed that amplitude correlations were more pronounced in the central and posterior channels while frontal channels showed a reduction of significant edges between recording sessions in at least 50% (Fig. S6a-b), 66% (Fig. 4c-d) of the subjects, there is no significant edge in at least 75% of the patients. Our analyses suggest that a prominent reduction of phase and amplitude coupling is present mainly between frontal, central, and inter- hemispheric channel groups in alpha and delta band, respectively. Of note, power increase associated with iRBD patients in delta rhythm during wakefulness does not reflect a more coherent brain activity.

### 3.4. Altered phase and amplitude couplings correlate with clinical and NPS scores

We asked whether altered phase and amplitude couplings correlate with Clinical score, NPS indices and mean SBR data. First, we computed Age-Adjusted Partial Pearson correlation coefficient of Clinical and NPS scores with wPLI in 10 Hz and oCC in 4 Hz (Fig. 5). In particular, we observed a direct positive correlation between global and inter-hemispheric wPLI values and visuo-spatial NPS score (*p* = 0.037 and *p =* 0.0094, respectively; BH corrected). On the other hand, the oCC showed an inverse correlation with MMSE and NPS data, and a positive correlation with PDSS-2. In particular, we observed a significant negative correlation between oCC and MMSE (*p* = 0.0031 – global and antero-posterior; *p =* 0.0052 – inter-hemispheric; *p* = 0.0043 – intra-hemispheric oCC configuration; BH corrected), verbal-memory (*p* = 0.0042 – global and intra-hemispheric; *p* = 0.0043 – inter-hemispheric; *p* = 0.0031 – antero-posterior oCC configuration; BH corrected) and executive index (*p* = 0.017 - global, *p* = 0.012 inter-hemispheric, *p =* 0.024 - intra-hemispheric and *p* = 0.0069 – antero-posterior oCC configuration; BH corrected).

**Fig.5:**
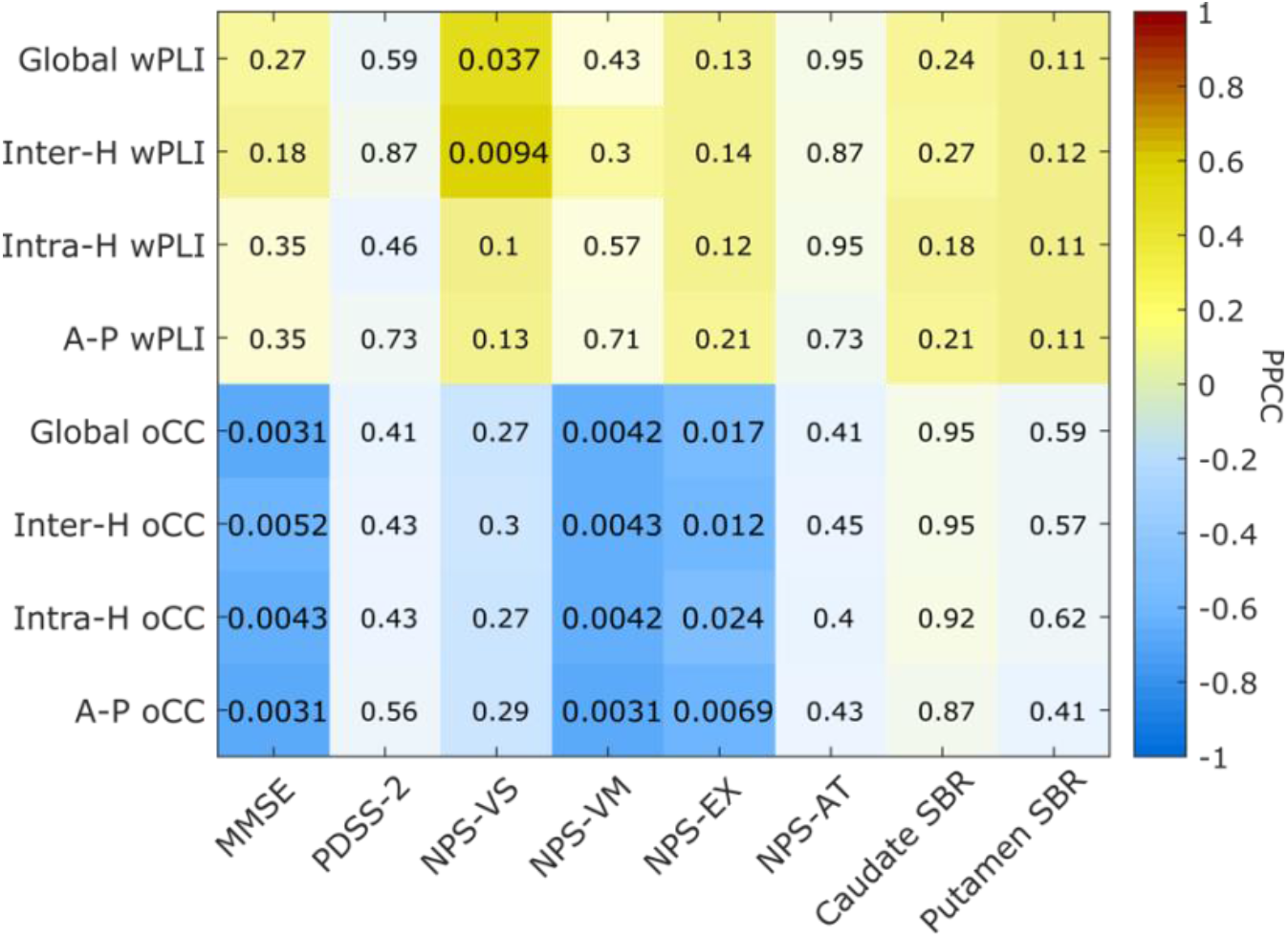
Altered phase and amplitude couplings correlate with clinical and NPS scores. Age-Adjusted Partial Pearson correlation coefficient with Benjiamini-Hochberg correction of wPLI (10 Hz) and oCC (4Hz) with clinical, cognitive and mean SBR data. We tested the alternative hypothesis that the correlation is not 0 with 95% of significance level. If the correlation coefficient is 1 there is a direct correlation, if it is 0 there is no correlation and if it is -1 there is an inverse correlation .In figure, we also reported the adjusted significant p-value. **Legend.** PPCC = Partial Pearson Correlation Coefficient; H = Hemispheric; A-P = Antero-Posterior; wPLI = weighted Phase Lag Index; oCC = orthogonalized Correlation Coefficient; MMSE= Mini-Mental State Examination; PDSS-2 = Parkinson’s Disease Sleep Scale; NPS-VS, NPS-VM, NPS-EX and NPS-AT = Neuropsychological visuo-spatial, verbal-memory, executive index and attention-mix, respectively; SBR= Specific to non-displaceable binding ratio.

## 4. Discussion

In this study for the first time, we longitudinally investigated iRBD patients by using serial HD-EEG, DAT-SPECT, clinical and NPS assessments. Patients were investigated at the time of iRBD diagnosis and then about two years later. First, we found a consistent shift of alpha power peak towards theta band of iRDB patients compared to age-matched healthy controls (Fig. 2). This is in agreement with previous literature data.^7–10,32,33^ At follow-up visit, the slowing of main alpha rhythm remained stable without clear worsening as disease progresses. However, by analyzing large-scale brain network coupling, we found a significant increase of the phase synchronization in the alpha (8-13 Hz) band in each channel groups (Fig. 3a-c, S3) from baseline to follow-up session. A large body of evidence showed the importance to study the phase-synchronization in prodromal and in early stage of neurodegenerative diseases.^11–13,34–37^

Hyper-synchrony in theta, alpha and beta bands was linked with disease progression,^11,13,37^ decreases at advanced stage of pathology.^13,36^ On the one hand, the increase of phase synchronization was associated with the activation of a brain compensatory mechanism to cognitive impairment.^11^ On the other hand, the hyper-synchronization was considered a sign of system malfunctioning that anticipate the breakdown of the network in the dementia stage.^13^ Altered phase synchronization may be observed in prodromal synucleinopathy as well. Indeed, putative hallmarks of cognitive decline and neurodegeneration include slowing of EEG alpha rhythms,^7^ alterations of global and network specific functional connectivity in both electrophysiological^38^ and resting-state functional MRI.^39–41^ In particular, in iRBD patients with cognitive impairment the power of alpha rhythm decreases and is replaced by an increase in delta/theta activity compared to age-matched healthy controls.^8^ In the present study, the increase of the phase synchronization in alpha band between baseline and follow-up was significantly and positively correlated with the neuropsychological scores (Fig. 5), especially in visuo-spatial functions. This finding supports the hypothesis that phase-synchronization increase can reflect the activation of a compensatory mechanism of the brain to counterbalance cognitive decline during neurodegeneration also in iRBD patients. It is also worth highlighting that the phase synchronization in alpha band showed a direct but not significant correlation (*p* = 0.11 – global, intra-hemispheric and antero-posterior; *p* = 0.12 – inter-hemispheric wPLI configuration; BH corrected) with nigro-striatal dopaminergic function (Fig. 5), in particular at putamen level, which is a known marker of neurodegeneration in iRBD patients and in synucleinopathy in general. Interestingly, we found that the phase-synchronization was enhanced in parietal and occipital lobes at follow-up and the hyper-synchronization was more pronounced in the right hemisphere (Fig. 4a-b, S5). This is in agreement with literature data showing that posterior brain regions are mainly involved in cognition in overt synucleinopathy like PD^42,43^ and DLB,^44^ and in prodromal synucleinopathy^39,45^ as well. Moreover, a recent study showed a significant asymmetry of brain degeneration in iRBD patients, with the left hemisphere being the most affected.^46^ Therefore, the increase in phase-synchronization in the right hemisphere might reflect the compensatory brain mechanism in the less affected hemisphere.

As a further result, we found a significant reduction of amplitude correlation in all lobes at follow-up compared with baseline (Fig. 3d-f, S4). In particular, this reduction was significant in the delta band (4 Hz) (Fig. 3f) and was inversely and significantly correlated with MMSE, verbal memory and executive functions (Fig. 5). RBD symptoms are known to affect large-scale global and between-region resting-state networks. Moment-to-moment correlation of EEG amplitude fluctuations shows spatial patterns similar to BOLD resting-state networks.^28^ Previous literature data found altered BOLD nigro-striatal, nigro-cortical and cortico-cortical activity iRBD patients.^39–41,47^ In particular, in agreement with our data a disruption of functional connectivity in the posterior brain regions has been associated with cognitive impairment in iRBD patients.^39^

Sleep disruption may play a role in cognitive function,^48^ thus we investigated whether large-scale functional couplings may be associated with sleep alteration. In our sample, we did not find any significant correlation between neither delta amplitude correlation or alpha phase synchronization and PDSS-2 scores (Fig. 5), thus we may speculate that the association between phase and amplitude couplings and cognitive function is independent from sleep disruption in our iRBD patients.

We can envisage some limitations of this study. First, the limited number of subjects involved limits the generalizability of our observations. This work constitutes a preliminary report of a larger project, the first of his kind, aiming at robustly investigating advanced electrophysiological markers in a longitudinal study of iRBD patients. Nonetheless, our results fit with current understanding of phase synchronization modifications in neurodegenerative disease and represent the first report of longitudinal electrophysiological modifications in prodromal stage of synucleinopathy. Second, volume conduction and signal mixing significantly inflate the phase synchronization and amplitude couplings. This effect is even larger for scalp recordings. For this reason, in the present work we adopted state-of-the-art metrics able to limit such bias. Moreover, a large fraction of clinical observations is still based on scalp level recordings and our results suggest that scalp large-scale network changes can be a useful biomarker in prodromal synucleinopathy. Finally, only two time-points cannot be used to infer accurate model prediction for disease progression. We are actively working towards increasing the number of subjects involved and the number of follow-up visits.

In conclusion, this study for the first time investigates the changes of phase-synchronization and amplitude correlation in iRBD subjects in two distinct time-points, by using HD-EEG, and correlating the data with longitudinal DAT-SPECT and cognitive assessment. Understanding the evolution over time of clinical, imaging and neurophysiological biomarker in iRBD could improve our understanding on the prodromal stages of synucleinopathy, possibly helping in identifying those patients that are more likely eligible for neuroprotective treatment. Our results suggest that global increase in alpha phase synchrony and the reduction of delta amplitude correlation might represent an electrophysiological correlate of the activation of a large-scale compensatory mechanism to balance cognitive decline.

## Acknowledgements

This work was developed within the framework of the DINOGMI Department of Excellence of MIUR 2018-2022 (legge 232 del 2016).

