## Supplementary Materials for "Phase and amplitude correlations change with disease progression in idiopathic Rapid eye-movement sleep behavior disorder patients"

### $^{123}$I-FP-CIT-SPECT image acquisition and reconstruction.

To perform SPECT images, we used a dual-head Millennium VG camera (GE Healthcare) equipped with low energy, high resolution, parallel-beam collimators. Scans have been acquired 180-240 minutes after intravenous administration of 185 MBq of $^{123}$I-FP-CIT (DaTSCAN, GE Healthcare, Little Chalfont, Buckinghamshire, UK), and lasted 40 minutes. We applied a "step-and-shoot" protocol with a radius rotation lower than 15 cm, and 120 projections evenly spaced over 360° were generated. Total counts were comprised between 2 and 3 million. We used an electronic zoom (zoom factor = 1.8) during data collecting phase to obtain an acquisition matrix's pixel size of 2.4 mm. Subsequently, a digital zoom was applied during the reconstruction phase. Resulting images were sampled by cubic voxels (2.33 mm). We processed projections by applying an OSEM algorithm (8 interactions, 10 subsets) that included a proback pair accounting for collimator blur and photon attenuation, and then post-filtering (3-D Gaussian filter with full-width at half maximum 58 mm). Photon attenuation was modelled with the approximation of a linear coefficient uniform inside the skull and equal to 0.11 $cm^{-1}$, and a 2D+ 1 approximation was applied in the simulation of the space simulation blur. Compensation for scatter was not performed.

### Basal Ganglia software functioning.

Basal Ganglia software is an automatic algorithm based on a high-definition, 3-dimensional striatal template derived from the Talairach atlas. In detail, an automated algorithm performs fine adjustments in positioning blurred templates to match radioactive counts, meanwhile, it locates occipital regions of interest (ROI) to perform background evaluation. Also, a partial volume effect (PVE) correction is performed during the uptake computation of the putamen, caudate and background.

| | NPS-VS | NPS-VM | NPS-EX | NPS-AT |
|---|---|---|---|---|
| Semantic verbal fluency | 0.43 | | | |
| Stroop Color | 0.54 | | | |
| TMT-A | -0.42 | | | |
| Clock completion test | 0.65 | | | |
| C.A. simple copy | 0.90 | | | |
| C.A. guiding landmarks | 0.94 | | | |
| RAVLT, immediate recall | | 0.81 | | |
| RAVLT, delayed recall | | 0.99 | | |
| Babcock story | | 0.85 | | |
| Stroop Color Word | | | 0.40 | |
| Corsi span | | | 1.02 | |
| TMT-B | | | -0.70 | |
| Symbol digit | | | 0.56 | |
| Phonemic verbal fluency | | | | 0.73 |
| Digit span | | | | 0.97 |
| **Variance explained (%)** | **63.6** | **13.2** | **10.1** | **7.9** |

***Table S1**: **Factor analysis results.** Factors and corresponding neuropsychological tests with their respective factor loading are shown. A conventional factor loading threshold of 0.4 was used. **Legend.** NPS-VS, NPS-VM, NPS-EX and NPS-AT = Neuropsychological visuo-spatial, verbal-memory, executive index and attention-mix, respectively; TMT= trail making test; C.A.= constructional apraxia; RAVLT= Rey Auditory Verbal Memory Test.*

| | AGE | MMSE | PDSS-2 | BDI-II | MDS-UPDRS-III | NPS-VS | NPS-VM | NPS-EX | NPS-AT | SBR Putamen | SBR Caudato |
|---|---|---|---|---|---|---|---|---|---|---|---|
| **S1** | 69 | 28 | 15,41[#] | 15 | 3 | 0,59 | -0,16 | 0,27 | 0,57 | 1.585 | 3.125 |
| | 71 | 27 | 2 | 12 | 3 | -0,44 | -1,97 | 0,19 | -0,42 | 0.705 | 2.575 |
| **S2** | 62 | 29 | 16,95[#] | 2 | 0 | 1,2 | -0,14 | 1,46 | 0,35 | 3.235 | 3.785 |
| | 64 | 30 | 7 | 3 | 0 | 1,23 | 0,44 | 1,45 | -0,34 | 2.685 | 3.125 |
| **S3**** | 77 | 24 | 10,70[#] | 7 | 2 | -1,87 | -2,33 | -2,69 | -0,59 | 1.75 | 2.85 |
| | 79 | 25 | 6 | 5 | 3 | -2,33 | -2,29 | -0,86 | -1,5 | 1.42 | 2.135 |
| **S4** | 72 | 27 | 38 | 19 | 8 | -0,028 | -0,21 | -0,49 | 0,45 | 2.795 | 3.73 |
| | 74 | 28 | 31 | 14 | 23 | -0,48 | 0,22 | -1,08 | -0,23 | 3.345 | 5.16 |
| **S5** | 71 | 29 | 17,33[#] | 12,6 | 0 | -2,31 | -0,58 | -0,097 | -1,78 | 2.745 | 3.56 |
| | 74 | 27 | 4 | 12 | 3 | -1,75 | -0,29 | 0,0096 | -1,62 | 3.015 | 3675 |
| **S6*** | 76 | 26 | 12,11[#] | 29,4 | 3 | -0,11 | -0,74 | -1,38 | 0,068 | 0.815 | 2.46 |
| | 78 | 25 | - | 10 | 15 | -0,17 | -0,83 | -1,83 | -1,03 | 0.815 | 2.19 |
| **S7** | 74 | 27 | 9 | 3 | 0 | -0,27 | 0,46 | -0,64 | -1,5 | 4.225 | 5.49 |
| | 76 | 29 | 4 | 3 | 0 | 0,11 | 0,56 | -0,29 | -1,52 | 2.96 | 3.51 |
| **S8** | 71 | 26 | 12,30[#] | 0 | 2 | -0,19 | -0,31 | 0,38 | 0,30 | 2.905 | 3.345 |
| | 74 | 27 | - | 6 | 6 | 0,25 | 0,14 | 1,11 | 1,08 | 2.85 | 3.015 |
| **S9** | 72 | 30 | 11 | 15 | 0 | 0,25 | 0,45 | 0,55 | 2,01 | 1.42 | 2.02 |
| | 73 | 28 | - | 25,2 | 10 | -1,75 | -0,045 | -1,25 | 1,67 | - | - |
| **S10** | 74 | 29 | 17,97[#] | 4 | 0 | 0,27 | 0,12 | 0,91 | 0,95 | 2.74 | 3.29 |
| | 77 | 28 | 20 | 9 | 0 | 0,091 | 0,64 | 0,49 | 0,91 | 2.905 | 3.62 |
| **S11*** | 67 | 28 | 20,29[#] | 16 | 4 | 0,25 | 1,2 | 0,15 | 0,59 | 1.035 | 1.145 |
| | 69 | 27 | 35 | 15 | 11 | 0,55 | 0,19 | 0,14 | 0,59 | 0.595 | 1.31 |
| **S12** | 60 | 29 | 24 | 17 | 0 | 0,62 | 0,77 | -0,55 | 0,56 | 3.29 | 3.455 |
| | 63 | 30 | 25 | 9 | 0 | 0,64 | 0,58 | 0,299 | 0,64 | 3.125 | 3.895 |
| **S13** | 69 | 29 | 17,73[#] | 10 | 0 | 0,25 | 0,95 | -0,44 | -0,22 | 2.35 | 2.63 |
| | 72 | 30 | 43 | 9 | 0 | 0,15 | 0,084 | -0,33 | -1,18 | 2.245 | 2.355 |
| **S14** | 77 | 30 | 17,45[#] | 16 | 1 | 0,099 | 0,97 | 0,63 | -0,092 | 2.85 | 3.29 |
| | 80 | 29 | 17 | 84 | 3 | 0,41 | 1,61 | 0,58 | 0,70 | 3.07 | 3.62 |
| **S15*** | 60 | 29 | 19,65[#] | 14 | 6 | 0,16 | -0,30 | -0,60 | -0,34 | 2.19 | 2.52 |
| | 62 | 29 | 38 | 25 | 16 | -0,57 | -1,18 | -0,91 | -0,57 | 2.575 | 3.325 |
| **S16** | 82 | 30 | 34 | 13 | 0 | 0,97 | 1,45 | 0,39 | 0,0019 | 2.905 | 4.115 |
| | 84 | 29 | 15 | 11 | 0 | 0,86 | 1,28 | 0,48 | 1,17 | - | - |
| **S17** | 77 | 29 | 24 | 6 | 0 | 0,49 | -0,36 | 0,63 | 0,68 | 2.3 | 2.905 |
| | 80 | 26 | 4 | 3 | 6 | 0,0040 | -1,26 | 0,096 | -0,44 | 1.97 | 3.29 |
| **S18** | 53 | 29 | 22 | 9 | 0 | 1,39 | -0,24 | 1,55 | -0,69 | 4.885 | 5.38 |
| | 55 | 30 | 11 | 3 | 0 | 1,45 | 1,10 | 1,67 | 0,86 | 4.28 | 4.445 |
| **mean ± SD** | 69,7 ± 7,5 | 28,2 ± 1,6 | 18,9 ± 7,7 | 11,6 ± 7,2 | 1,6 ± 2,4 | 0,098 ± 0,9 | 0,056 ± 0,9 | 0,002 ± 1,0 | 0,07 ± 0,9 | 3,3 ± 1,1 | 2,6 ± 1,0 |
| | 71.7 ± 7.5 | 28,0 ± 1,6 | 17,5 ± 13,9 | 11,0 ± 7,3 | 5,5 ± 6,8 | -0,098 ± 1,0 | -0,056 ± 1,1 | -0,002 ± 0,9 | -0,07 ± 1,0 | 3,2 ± 0,9 | 2,4 ± 1,0 |
| ***p*** | n.a. | 0.66 | 0,29 | 0,58 | 0,15 | 0,46 | 0,94 | 0,79 | 0,86 | 0.91 | 0.88 |

***Table S2: Cognitive decline effects are not apparent with disease progression****. On the rows, for each subjects, we observed the baseline (up) and the follow-up (bottom) values for each index. The two last row showed the mean ± standard deviation for both baseline and follow-up, and the p-value of the Wilcoxon test computed between baseline and follow-up values, respectively. * indicates the RBD patients which are evolved in PD at follow-up, while ** indicates the patients which are evolved in DLB. The PDSS-2 is estimated using an alternative method in baseline, marked with #.* ***Legend.*** *BDI-II= Beck Depression Inventory II; MDS-UPDRS-III= Movement Disorder Society-Unified Parkinson's Disease Rating scale, motor section; MMSE= Mini-Mental State Examination; NPS-VS, NPS-VM, NPS-EX and NPS-AT = Neuropsychological visuo-spatial, verbal-memory, executive index and attention-mix, respectively; SBR= Specific to non-displaceable binding ratio. PDSS-2 = Parkinson's Disease Sleep Scale; SD = Standard Deviation.*

| | Delta Rhythm | Theta Rhythm | Alpha Rhythm | Beta Rhythm |
|---|---|---|---|---|
| **Recording sessions** | p = 0.67 | p = 0.70 | p = 0.71 | p = 0.61 |
| **Electrode groups** | p = 0.068 | p = 0.074 | p = 0.018 | p = 0.27 |
| **Sessions x Groups** | p = 0.34 | p = 0.34 | p = 0.28 | p = 0.32 |

***Table S3: Repeated Measure ANOVA two-ways over Relative Power values.*** *We compared the mean difference between the recording session in each brain rhythm (delta, theta, alpha and beta).*

| | $\alpha\ (1/f^{\alpha})$ |
|---|---|
| **Recording sessions** | p = 0.64 |
| **Electrode groups** | p = 0.52 |
| **Sessions x Groups** | p = 0.064 |

***Table S4: Repeated Measure ANOVA two-ways over slope alpha values.*** *We compared the mean difference between the recording sessions.*

| | Delta Rhythm | Theta Rhythm | Alpha Rhythm | Beta Rhythm |
|---|---|---|---|---|
| **Recording sessions** | p = 0.68 | p = 0.71 | p = 0.72 | p = 0.63 |
| **Electrode groups** | p = 0.011 | p = 0.011 | p = 0.0017 | p = 0.056 |
| **Sessions x Groups** | p = 0.31 | p = 0.29 | p = 0.26 | p = 0.29 |

***Table S5: Repeated Measure ANOVA two-ways over Absolute Power values.*** *We compared the mean difference between the recording sessions.*

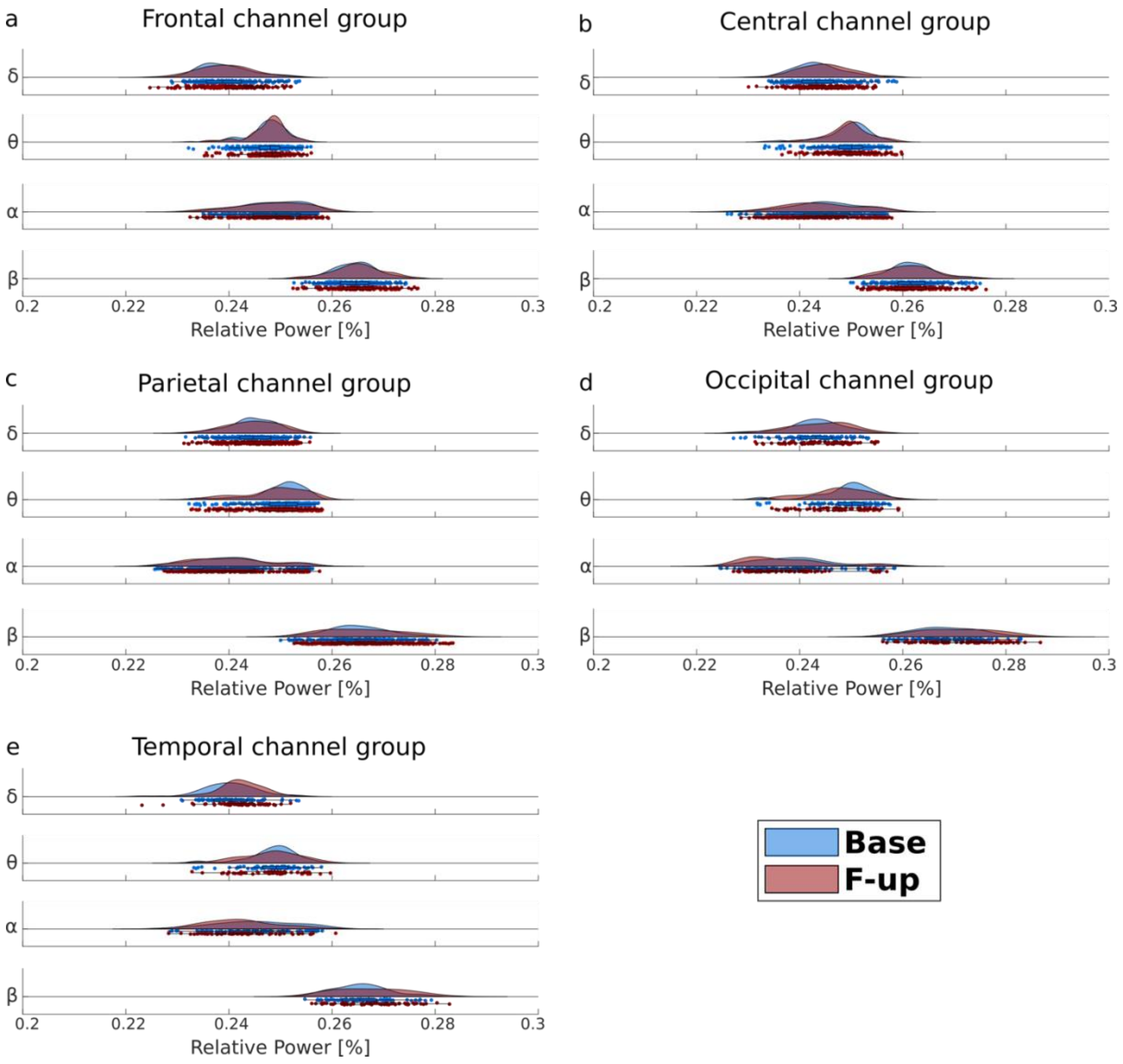


***Figure S1**: **Disease progression effects on amplitude distributions**. Raincloud plots of relative power as a function of frequencies for all 5 channel groups: frontal (a), central (b), parietal (c), occipital (d) and temporal (e). Dots represent relative power of each single channel pooled across subjects at baseline (blue) and at follow-up (red). **Legend.** Base = baseline; F-up = follow-up.*

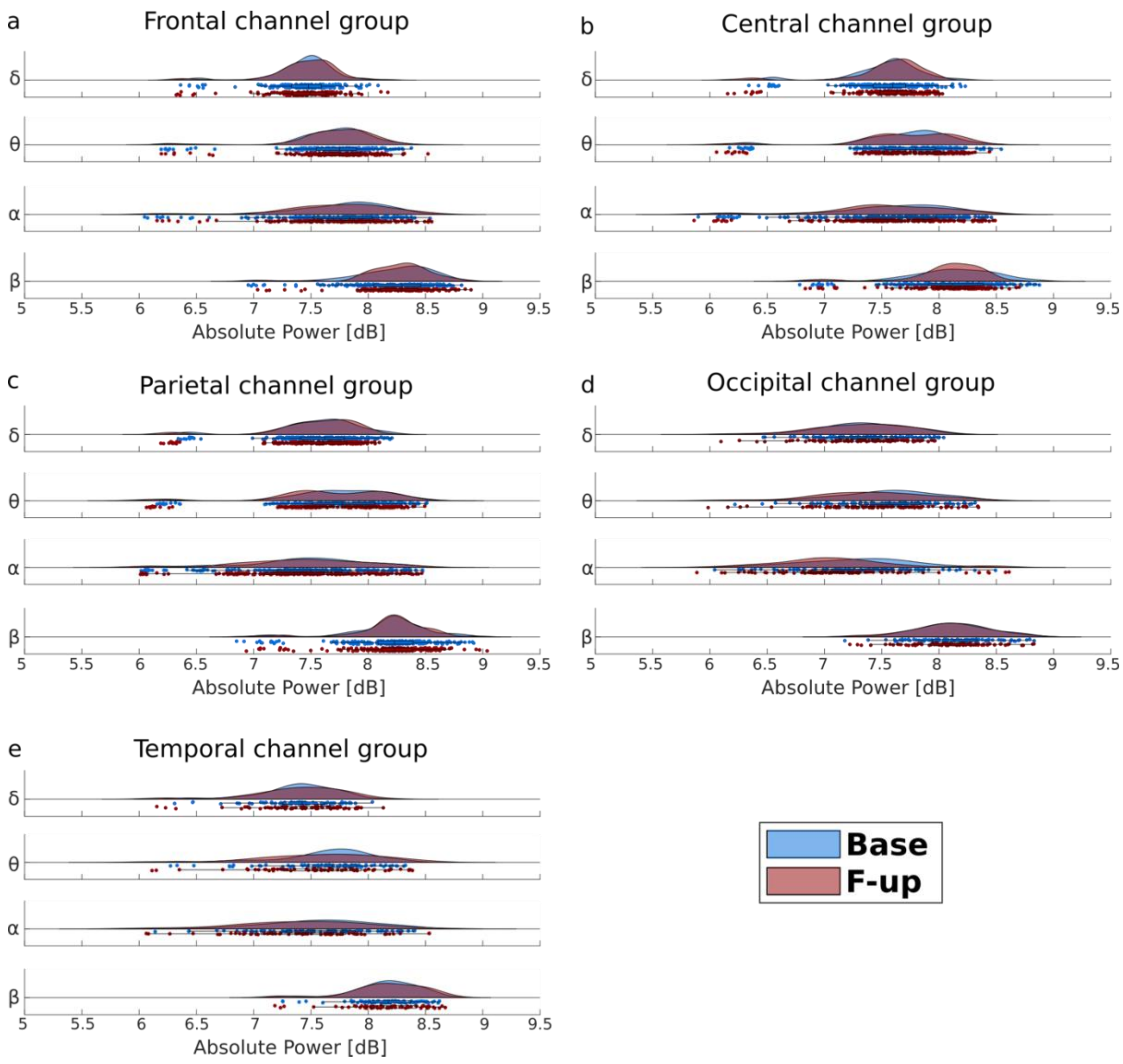


***Figure S2: Disease progression effects on amplitude distributions.*** *Raincloud plots of absolute power as a function of frequencies for all 5 channel groups: frontal (a), central (b), parietal (c), occipital (d) and temporal (e). Dots represent absolute power of each single channel pooled across subjects at baseline (blue) and at follow-up (red).* ***Legend.*** *Base = baseline; F-up = follow-up*

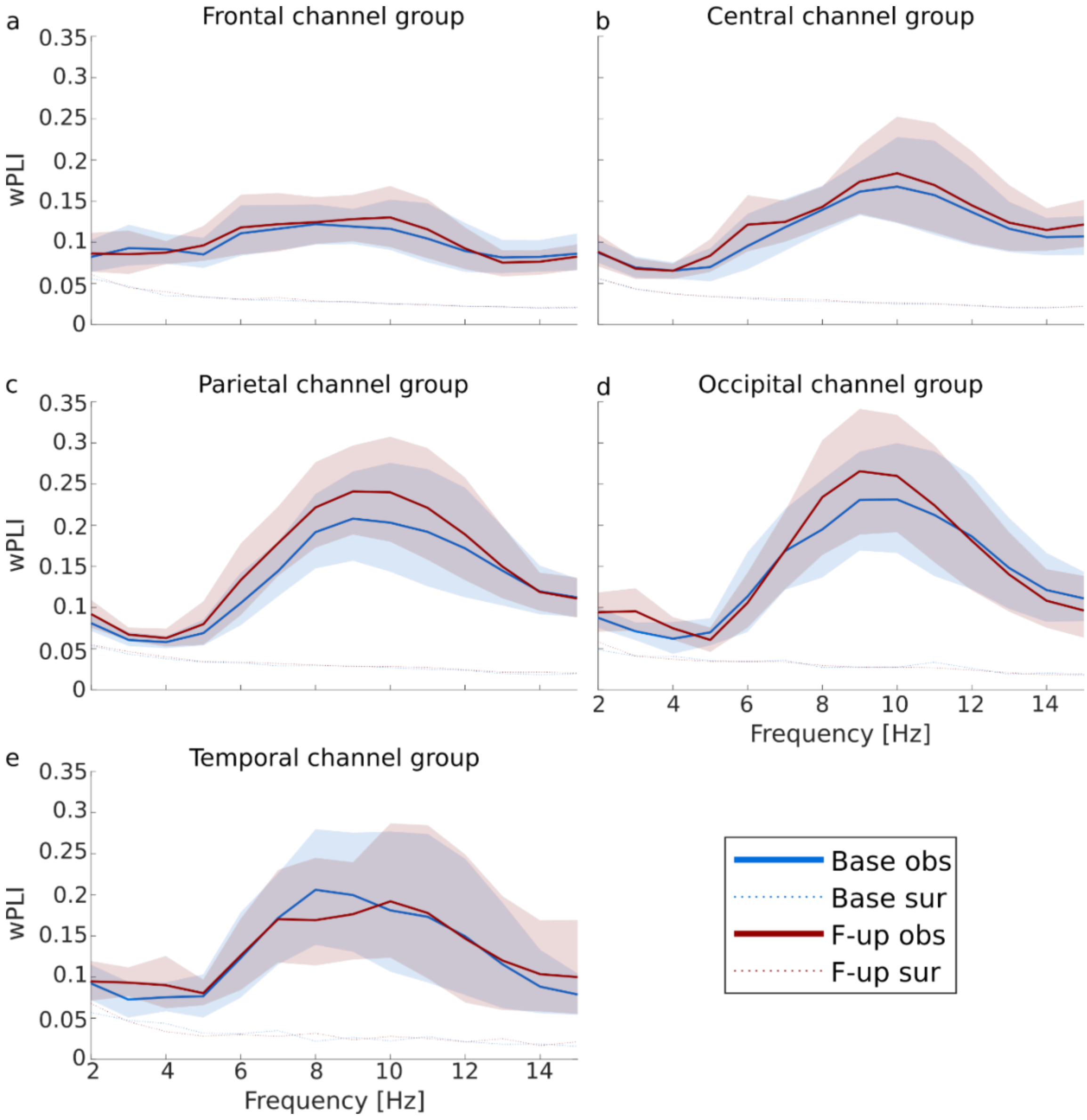


***Figure S3: Disease progression enhance the phase-synchronization in alpha band.*** *Strength of population averaged weighted Phase Lag-Index (wPLI) at baseline (blue) and at follow-up (red) in each channel group: (a) frontal, (b) central (c) parietal, (d) occipital and (e) temporal. Shaded areas represent confidence interval around population mean (1000 bootstraps). Dashed lines represent surrogate average.* ***Legend.*** *wPLI = weighted Phase Lag Index; Base = baseline; F-up = follow-up.*

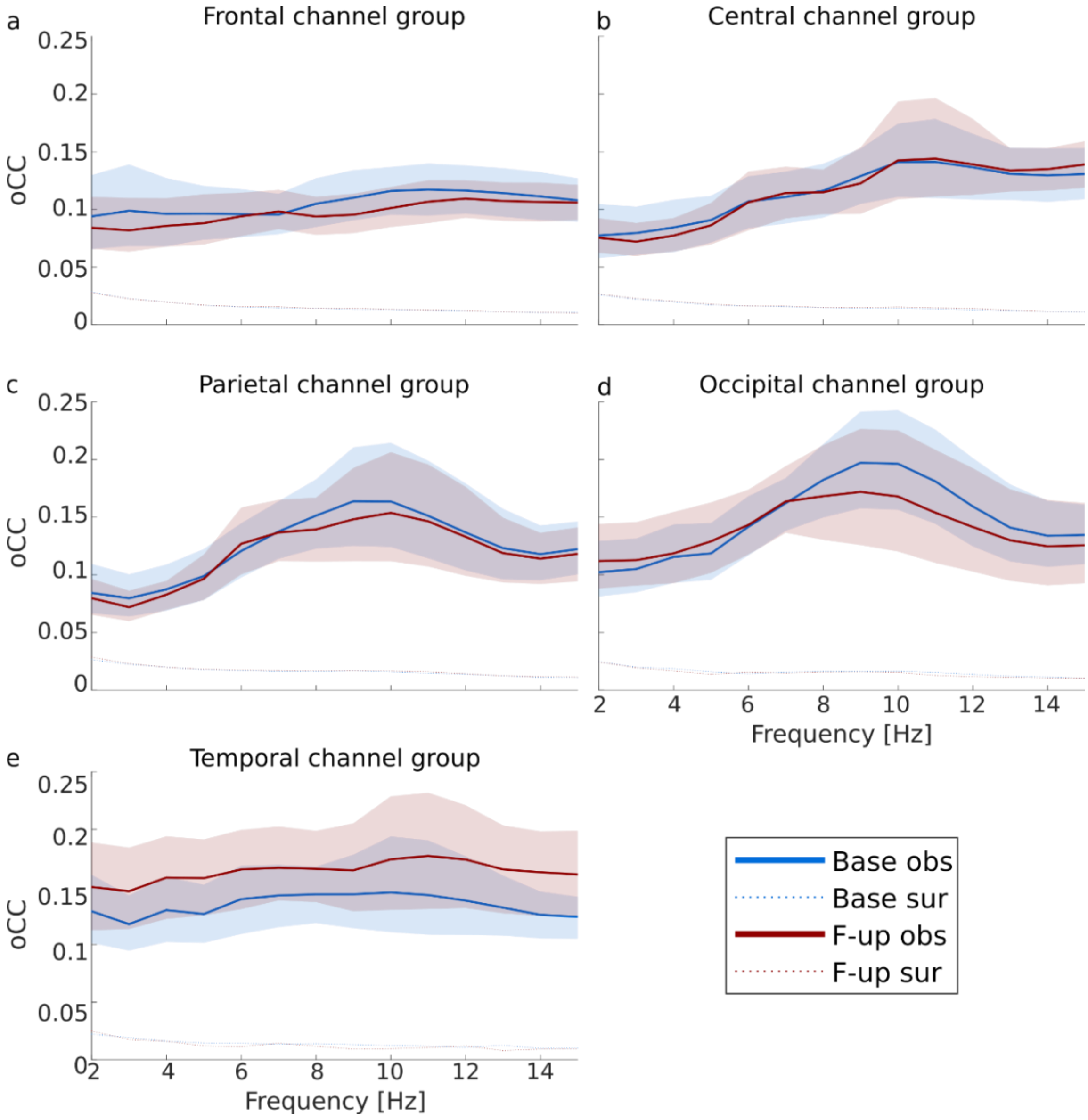


***Figure S4: Disease progression enhance the amplitude correlation in delta band.*** *Strength of population averaged Orthogonalized Correlation Coefficient (oCC) at baseline (blue) and at follow-up (red) in each channel group: (a) frontal, (b) central (c) parietal, (d) occipital and (e) temporal. Shaded areas represent confidence interval around population mean (1000 bootstraps). Dashed lines represent surrogate average.* ***Legend.*** *oCC = orthogonalized Correlation Coefficient; Base = baseline; F-up = follow-up.*

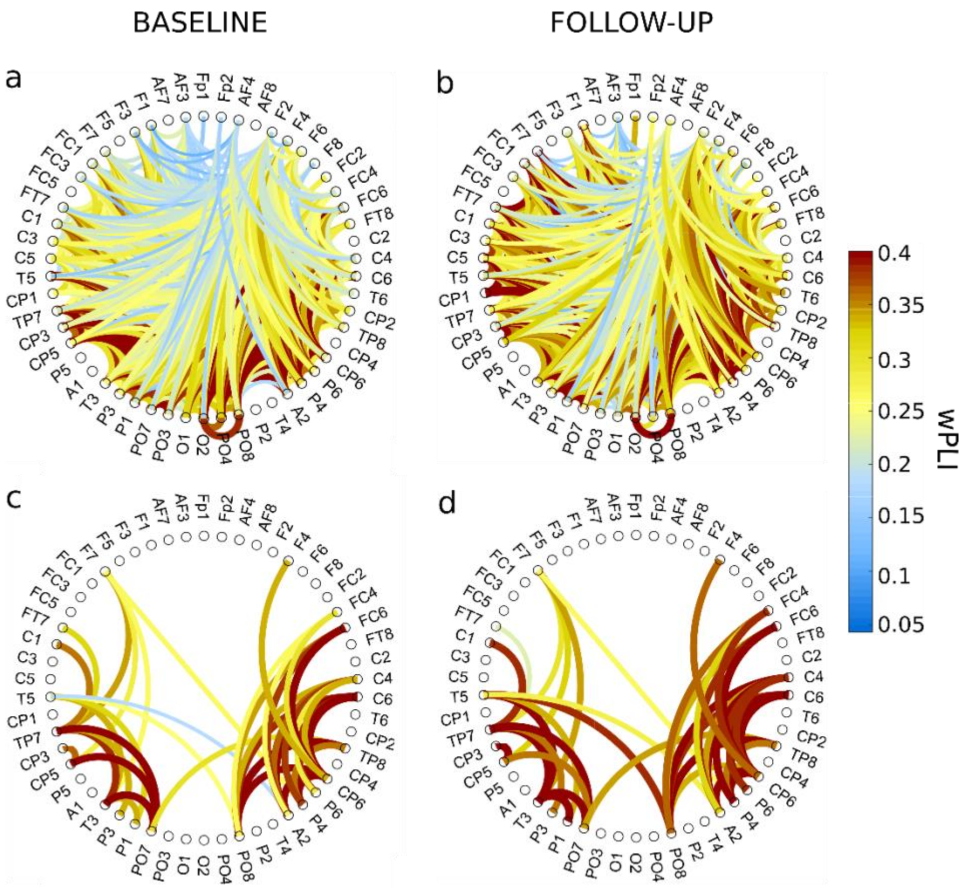


***Figure S5: Phase-synchronization increases in alpha band.*** *Graph of population averaged wPLI in alpha band (10Hz at baseline (left) and at follow-up (right) for significant edges (p<0.05) in at least 50% (a,b) and 75% (c,d) of the patients.* ***Legend:*** *wPLI = weighted Phase Lag Index*

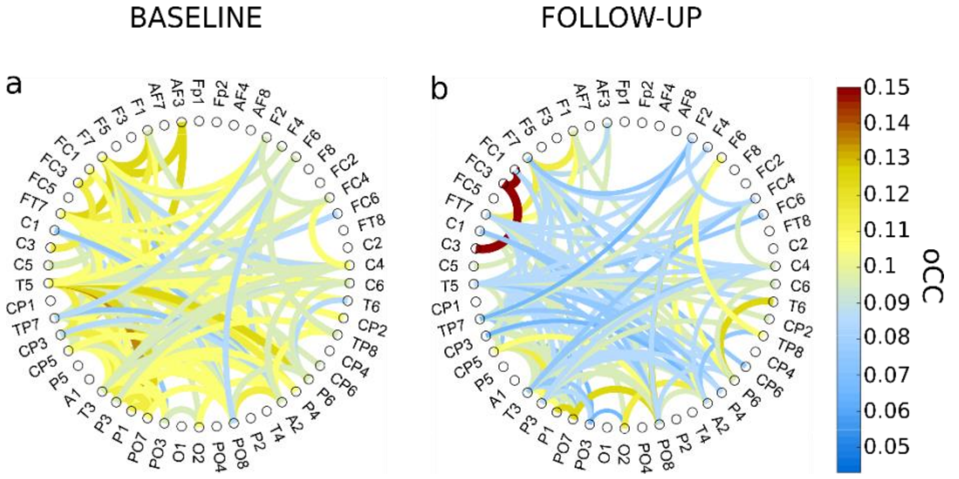


***Figure S6: Amplitude correlation decreases in delta band.*** *Graph of population averaged oCC in delta band (4Hz) at baseline (a) and at follow-up (b) for significant edges (p<0.05) in at least 50% of the patients.* ***Legend:*** *oCC = orthogonalized Correlation Coefficient*
